# Traveling waves in the monkey frontoparietal network predict recent reward memory

**DOI:** 10.1101/2022.02.03.478583

**Authors:** E. Zabeh, N.C. Foley, J. Jacobs, J.P. Gottlieb

## Abstract

Brain function depends on neural communication, but the mechanisms of this communication are not well understood. Recent studies suggest that one form of neural communication is through traveling waves (TWs)—patterns of neural oscillations that propagate within and between areas. We show that TWs appear robustly in microarray recordings in monkey frontal and parietal cortex and encode memory for recent rewards. While making saccades to obtain probabilistic rewards, monkeys were sensitive to the (statistically irrelevant) prior reward, which is consistent with behavioral biases produced by reward history. TWs in frontal and parietal areas were stronger in trials following a prior reward versus a lack of reward and, in the frontal lobe, correlated with the monkeys’ sensitivity to the prior reward. The findings suggest that neural communication across fronto-parietal areas, reflected by TWs, maintains default reward memories, while communication within the frontal lobe mediates the read out of the memories for prospective expectations.

## Introduction

Neural oscillations have long been proposed to regulate communication among neuronal ensembles within and across different brain structures (Fell and Axmacher, 2011; Fries, 2005). Traditionally, neural oscillations are most often interpreted as showing that large groups of neurons all experience so-called zero-lag synchrony, in which network activity fluctuates rhythmically with the same timing across cells. However, with the advent of simultaneous multichannel recording technologies, mounting evidence shows that many oscillations are in fact traveling waves (TWs)—oscillatory patterns of activity that propagate across neural tissue at speeds consistent with axonal conduction velocities.

TWs have been found in multichannel recordings of local field potentials across multiple animal species, frequencies, brain states, and brain systems, suggesting that they are a widespread feature of neural activity. Importantly, in addition to being found during sleep (Dickey et al., 2021; Massimini et al., 2004; Muller et al., 2016), TWs are prominent during active behaviors in animals – including in the hippocampus of rodents during navigation (Lubenov and Siapas, 2009; Patel et al., 2012), and visual regions of monkeys during perception (Davis et al., 2020, 2021) – as well as in humans performing visual (Alamia and VanRullen, 2019; Alamia et al., 2020; Halgren et al., 2019; Pang et al., 2020) and memory tasks (Kleen et al., 2021; Zhang and Jacobs, 2015; Zhang et al., 2018). In some of these studies, specific TW properties (e.g., strength or direction) correlate with accuracy and reaction times (Balasubramanian et al., 2020; Davis et al., 2020; Zhang et al., 2018) suggesting that TWs are functionally significant in linking neural activity with behavior.

The local field potentials that form TWs reflect the mean population activity of the neurons underneath a local electrode (Buzsáki et al., 2012; Katzner et al., 2009). Thus, a propagating TW indicates that there is a spatiotemporal pattern of neural activity— a wave of neuronal spiking—that is moving in a particular direction across a population of cells (Sato et al., 2012; Swadlow and Alonso, 2009; Takahashi et al., 2015). Because TWs activate ensembles of cells in succession, they may help integrate information coded in distinct populations of cells, such as linking visual information across different retinotopic locations or across a saccade (Davis et al., 2020; Zanos et al., 2015). However, much remains unknown about whether, and how, TWs encode the integration of information in time, as a potential memory mechanism.

Here we studied this question by examining the role of TWs in reward memory. Learning from past punishments and rewards is a cornerstone of behavior, and often occurs by default even based on statistically irrelevant recent events (in so-called “model-free” fashion; (Akrami et al., 2018; Abrahamyan et al., 2016; Lee et al., 2012)). Frontal and parietal areas are strong candidates for mediating this process as they contain individual neurons that encode reward expectations and have sustained cross-trial activity (Foley et al., 2020; Genovesio et al., 2014; Lee et al., 2012; Mansouri et al., 2006; Scott et al., 2017). To examine if TWs play a role in this process, we obtained microelectrode recordings from the dorsolateral prefrontal cortex (dlPFC) and parietal area 7A while monkeys made saccades to obtain probabilistic rewards that were uncorrelated across consecutive trials. We show that oscillatory potentials in the beta frequency band form reliable TWs that encode recent reward history. TWs at the start of a trial were stronger if the previous trial produced a reward relative to a lack of reward. The enhancement of TWs by a prior reward was found in both areas, but the influence of the reward on the monkeys’ expectations was specifically reflected by TWs in the dlPFC. Thus, TWs convey distinct reward information relative to individual neurons and local LFP oscillations and reflect distinct contributions of the frontal and parietal lobes to the maintenance and use of recent reward memories.

## Results

### Monkeys are sensitive to irrelevant prior rewards

Two monkeys performed a visually guided saccade task in which they obtained probabilistic rewards predicted by visual cues (Foley et al., 2020). In each trial, the monkeys saw a visual cue specifying the trial’s expected value (EV; combination of magnitude and probability; Fig. 1A). After maintaining fixation for an additional delay period, the monkeys made a saccade to a target to obtain the reward (Fig. 1A). The cues and target locations were independently randomized among two possible locations in the right or left visual hemifields, so that the initial cue signaled reward expectations but not the saccade motor plan. The EV of each trial was selected randomly and independently of the previous trial reward.

**Fig1.**
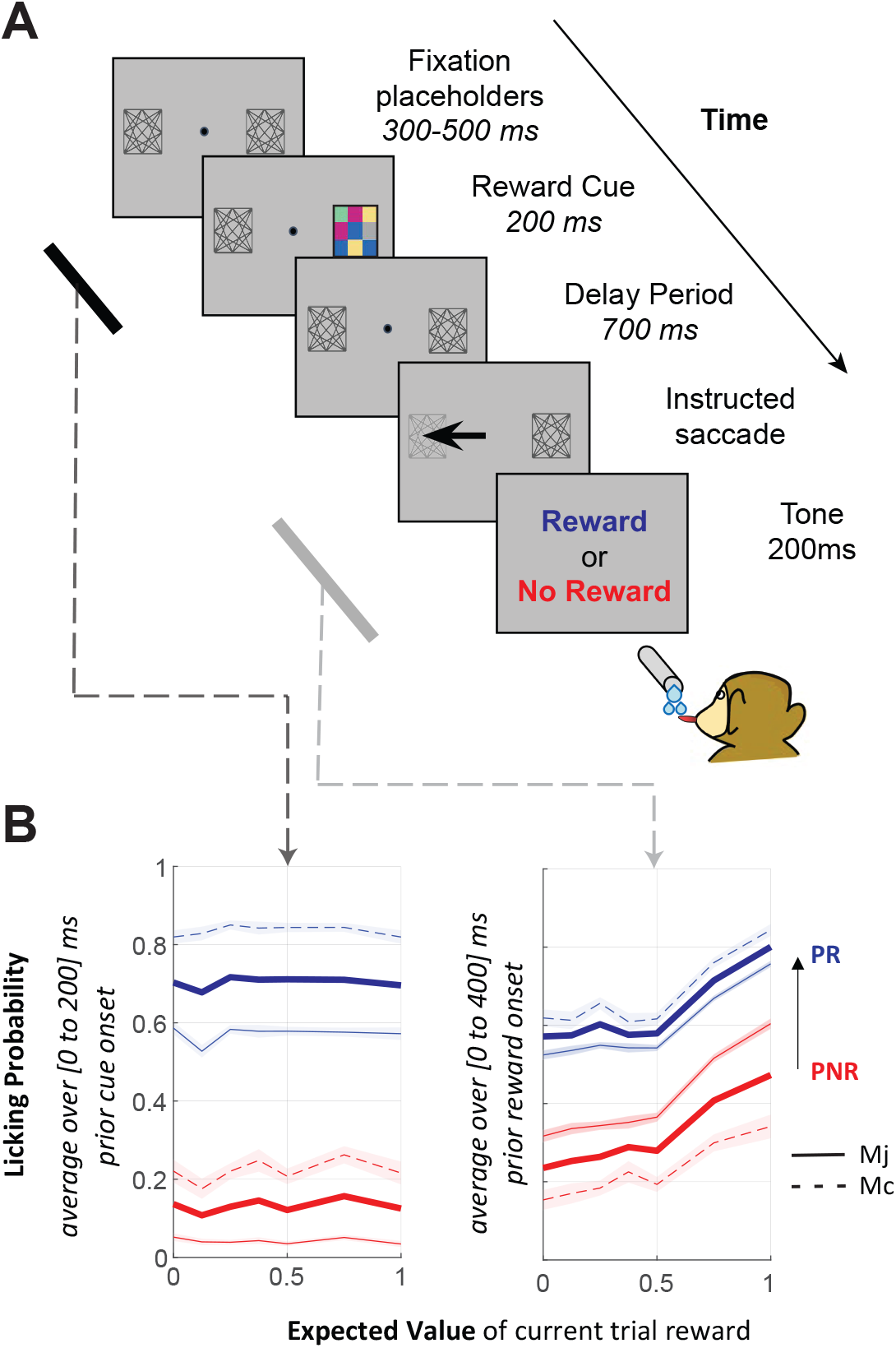
Task and behavior. **(A) Trial events**. On each trial, after achieving fixation, the monkeys saw a reward cue (colored checkerboard) indicating the trial’s EV. After a 700 ms delay period, a saccade target appeared and, after making the required saccade, the monkeys received the outcome (reward or lack of reward), according to the cued probability. Two gray placeholders were continuously present on the screen, marking the possible locations of the cue and target (which were randomized independently across trials). **(B) The probability of anticipatory licking depends on EV and the prior trial reward**. The traces show the licking probability as a function of EV during two task epochs: the fixation period (left, the 200 ms epoch ending at reward cue onset) and the delay period before reward delivery (right: 800 – 1200 ms after cue onset). Blue traces indicate trials that followed a rewarded trial (prior reward, PR) and red indicate trials that followed a lack of reward (prior no reward, PNR). Each trace shows the mean and SEM across all correct trials for each monkey (Mj, solid, Mc, dashed) and across both monkeys. Regression analysis showed that the effect of the prior trial outcome was significant in both monkeys in both the pre-cue and delay epochs (all p < 0.001)

We recorded the monkeys’ anticipatory licking (prior to reward delivery) as an index of their reward expectations. Licking rates increased with the EV signaled by the cues, confirming that the monkeys were familiar with and attentive to the reward cues (Fig. 1B, right). Surprisingly, licking also correlated with prior trial reward, despite the fact that this reward was not predictive of the current trial expectations (Fig. 1B, red vs blue traces). Licking was more vigorous if the prior trial ended with a reward (PR) versus a lack of reward (PNR), and this difference was seen during the period of fixation preceding the cue (Fig. 1B, left) as well as during the post-cue delay period (Fig. 1B, right). The prior trial effect did not merely reflect consumption of the prior reward, because licking ceased during the inter-trial interval and resumed at the start of the next trial (Foley et al., 2020). Licking rates were correlated with the rewards on the previous trial but not with those further removed in the past (2 trials back, Mc: r=0.003 p=0.43; Mj: r = 0.001 p = 0.46). Thus, consistent with prior reports (Lak et al., 2020) the monkeys’ reward expectations were sensitive to the recent reward history even though this history was irrelevant for behavior, consistent with previous findings in humans and monkeys (Abrahamyan et al., 2016; Genovesio et al., 2014).

### Oscillatory activity in the dlPFC and PPC shows traveling waves

We recorded local field potentials (LFPs) using electrode grids implanted in the dorsolateral prefrontal cortex (dlPFC) and posterior parietal cortex (PPC; area 7a) in each monkey (Fig. 2A). The LFPs at many individual electrodes showed prominent oscillatory activity in the beta-band (∼15 Hz). In addition, across individual electrodes, the oscillations had phase shifts with a consistent spatial gradient, indicating the presence of traveling waves, or plane waves, propagating in consistent directions. Fig. 2B shows an example of this phenomenon in one trial recorded from the PPC of monkey Mc. The LFP signals showed a similar oscillation waveform on neighboring electrodes (Fig. 2B, top two rows) and the individual oscillations had a systematic phase shift across adjacent electrodes (Fig. 2B, bottom row, electrodes 1-5). The phase shift was consistently oriented across the array, indicating that the oscillation propagated in an anterior-to-posterior direction across the grid of electrodes (Fig 2C).

**Fig2.**
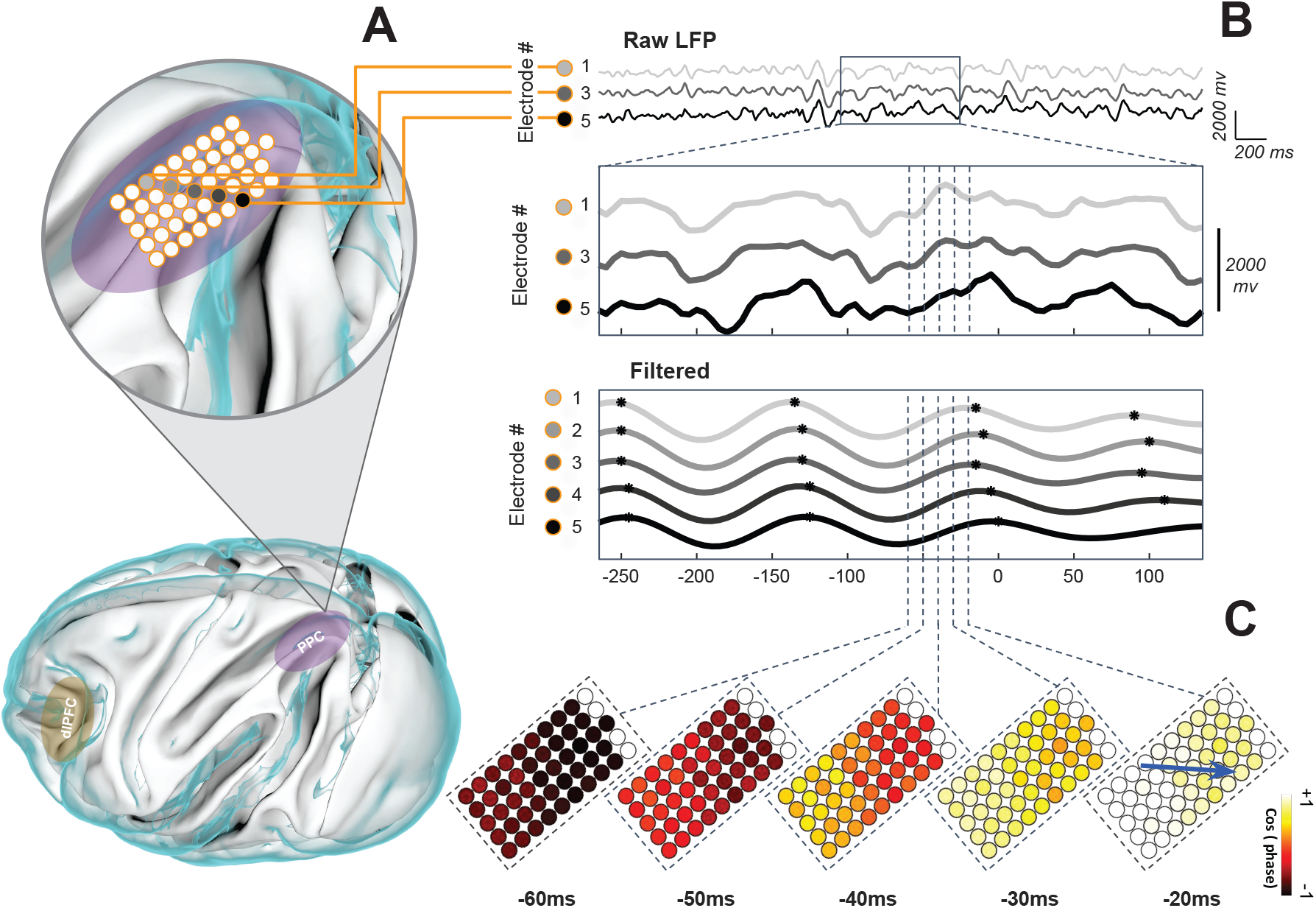
Spatial organization of oscillations in frontoparietal network. **(A) Locations of the implanted arrays**. The bottom panel shows the approximate locations of the PPC (purple) and dlPFC (yellow) recording areas approximated based on stereotactic coordinates and intra-operative photographs which was projected on a canonical 3D lateral view of the cortex, exported from diffusion tensor MRI atlas of the rhesus macaque brain (Bakker et al., 2015; Calabrese et al., 2015). The top panel shows a closeup of the PPC array from monkey Mc. The highlighted electrodes are those corresponding to the traces in B. (**B) A traveling wave in a representative trial from the electrodes highlighted in Fig. 2A**. The top panel shows the unfiltered LFP signals from every other electrode in the group, and the middle panel is an expanded view of a 400 ms time window of the same LFPs aligned on cue presentation onset (0 ms). The bottom panel shows the signals at all 5 electrodes filtered at 14 Hz (Bandwidth 1.5Hz). The oscillation peaks (black dots) occur at successively later times for electrodes 1-5. **(C) The traveling wave across the entire array**. The color shows the cosine of relative phase for the 14 Hz LFPs across the array at 10 ms intervals between 60 and 20 ms before cue onset. The blue arrow indicates the direction of the traveling wave defined as the direction of the maximum phase gradient at time -20 ms.

To test for traveling waves throughout the dataset, we measured the strength and direction of the spatial gradients of the LFP oscillations using circular–linear statistics (Fisher, 1993). We computed these measures for each frequency in the range of 2 to 50 Hz, and in the time interval from 1.2 seconds before to 2 seconds after cue onset (see *Methods*; Supplemental Movie S1, S2). Using this approach, we extracted the Phase Gradient Directionality (PGD) index, which provides a measure of the traveling wave strength (WS) across the grid at each frequency and timepoint during the task

This analysis revealed prominent TWs in all four recording arrays. Although TWs could appear at multiple frequencies, TWs were strongest in the alpha-beta band frequency range (10–30 Hz), with the peak frequency being slightly higher in the dlPFC (Mc 19.1Hz; Mj 13.6Hz) relative to the PPC (15.6 Hz vs 11.5 Hz; rank-sum test between PFC-PPC pairs, all p<0.001; Fig. 3A-C). Although we detected TWs at other frequencies (Fig. S2), across the arrays the frequency showing the strongest traveling waves was correlated with the frequency showing the strongest LFP oscillations (Fig. 3C; r^2^ = 0.54, p<0.01), suggesting that TWs result from propagation of the LFP fluctuations in the dominant frequency band. The speed of TW propagation was slightly higher in the dlPFC relative to the PPC (Wilcoxon rank-sum test: * P<0.001) but in both areas was in the range of 0.1–0.6 meters/second (Fig. 3D), matching the conduction velocity in unmyelinated axons (Girard et al., 2001; González-Burgos et al., 2000)

**Fig3.**
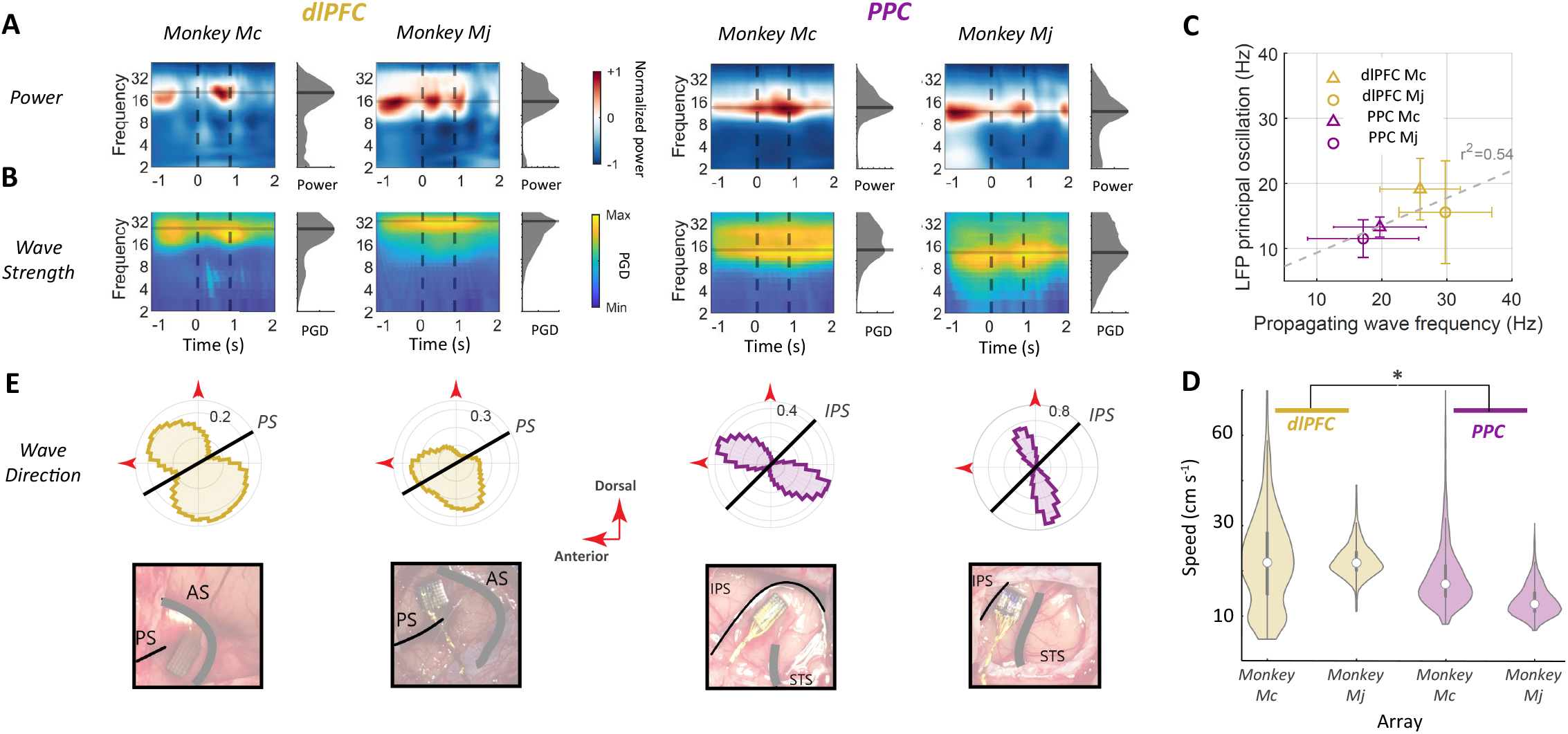
Properties of traveling waves. **(A) LFP power peaks in the beta band in each monkey and area**. Wavelet spectrograms were computed independently for each electrode and trial and averaged for each array (left: dlPFC; right: PPC). Frequency has log scaling and time is in milliseconds relative to cue onset. Power is normalized based on the maximum power pixel value in each array. The black dashed lines show, respectively, cue onset and target onset. The gray plot shows the average LFP power as a function of frequencies and the shaded horizontal lines indicate the peak frequency. Peak LFP power (red) was in the beta-band during the pre-cue, and delay epochs. **(B) Wave strength peaks in the beta band**. Wave strength was calculated on each trial and averaged across trials for filtered LFP signals on each trial, with center frequencies from 2 to 50 Hz. The gray marginal histogram shows the time-averaged wave strength as a function of frequency. The color codes are calculated independently for each array. **(C) Peak TW frequency correlates with peak LFP power frequency**. The frequency showing the strongest traveling wave correlated with that showing the highest LFP power (r2 = 0.54, p<0.01). Each point shows the mean and standard deviation of the peak frequency for LFP power and wave strength in individual trials for each array. **(D) TW speeds are biologically plausible**. Violin plots showing the across trial distributions of peak beta TW speeds averaged over the interval from -1.2s to 2 s relative to the cue. The white dots show medians, and the whiskers show standard deviations. The propagation speed of the beta wave is consistent with the conduction velocity in unmyelinated axons [10–60 cm per s ](Girard et al., 2001) and the PFC wave is slightly faster than PPC (rank-sum test: * P<0.001; across both monkeys). **(E) TWs propagate along consistent axes**. The circular histograms show the distribution of peak beta frequencies (PFC: Mc 25.7Hz, Mj 29.7Hz; PPC: Mc 19.7Hz, Mj 17.1Hz; Fig3.B). TW directions across all recording trials from -1.2 to 2 s relative to cue onset. The histograms are referenced to the dorsal and anterior directions and the black lines denote the axes of the principal sulcus (PS) in PFC and the intraparietal sulcus (IPS) in the PPC. The bottom row shows intraoperative photographs corresponding to each distribution, showing the locations of the arrays relative to the PS and IPS and the adjacent sulci (arcuate sulcus, AS, for PFC and the superior temporal sulcus, STS, for PPC).

The directions of the TWs were not random but tended to be approximately perpendicular to the nearby sulci, which is the principal sulcus in the dlPFC and the intraparietal sulcus in the PPC (Fig. 3E). TWs were equally likely to occur in both directions along these perpendicular axes, resulting in bimodal distributions with peaks in anterior-dorsal and posterior-ventral directions (Omnibus circular non-uniformity test: p < 0.001 for all arrays). The directional distributions were more narrowly tuned, suggesting that propagation directions were more consistent in the PPC relative to the PFC (the mean angular distance from the bimodal axis was, in Mc, PPC: 26.93±0.03° vs dlPFC: 37.88±0.04°, p <0.01; and in Mj; PPC: 26.64±0.03° vs dlPFC: 43.58±0.04°, p < 0.01, circular Kuiper test).

### Traveling waves correlate with prior rewards

To probe the functional relevance of traveling waves, we next analyzed the relation between their properties and behavior. We found a significant association between the strength of the TWs and the reward that the monkey received on the previous trial. Fig. 4A illustrates this result in two representative trials from the PPC array in monkey Mj. One trial, shown in the top panel, followed a prior reward (PR). In this trial, during the initial fixation period before the reward cue, the phase gradients at different electrodes were organized in the same direction, resulting in a high wave strength (0.42) indicating a traveling wave propagating towards 180 degrees. The second trial, shown in the bottom panel in Fig. 4A, followed a prior no-reward (PNR). In this trial, phase gradient directions were randomly organized, resulting in low wave strength (0.12) that suggested the absence of a traveling wave.

**Fig4.**
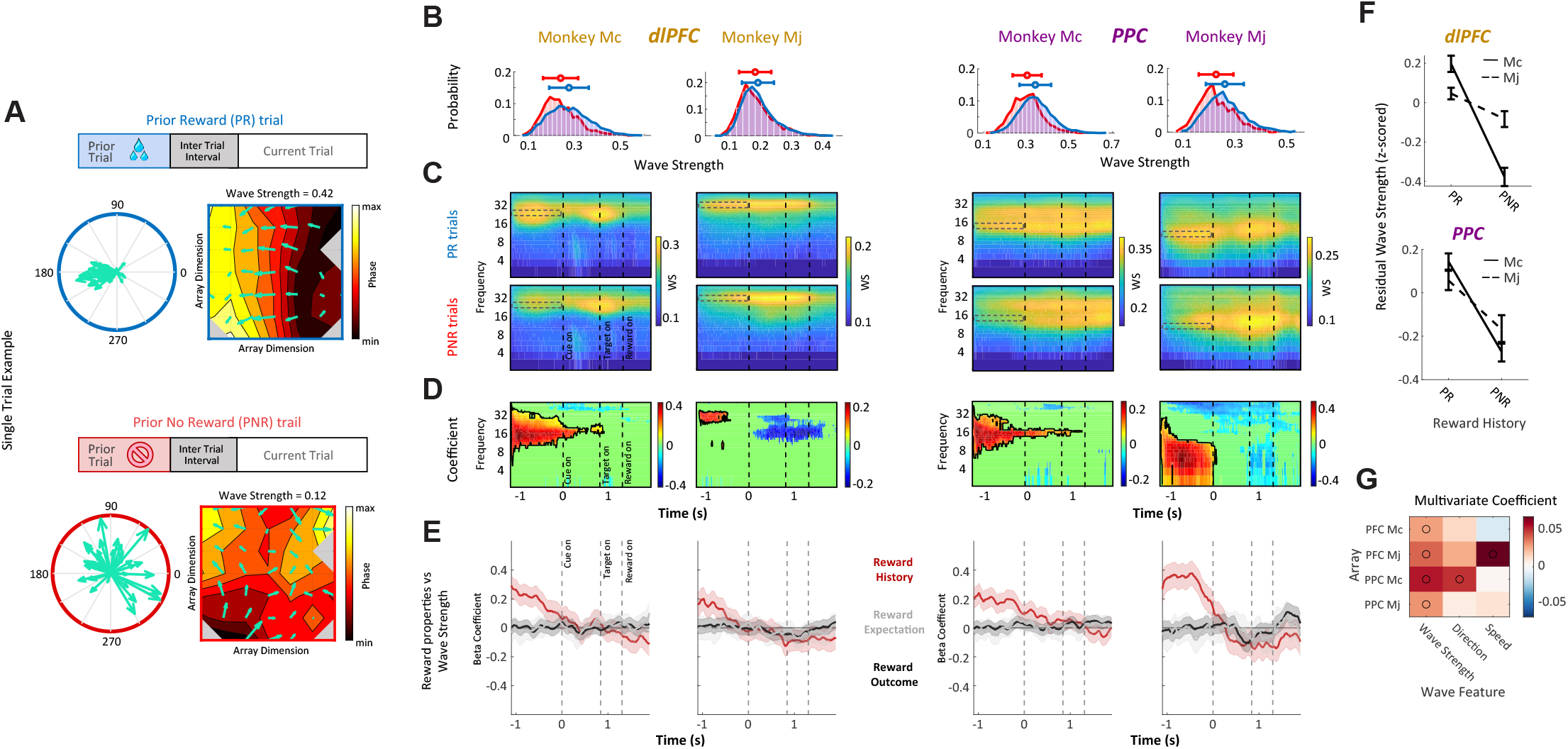
TW strength depends on the prior trial reward. **(A) Two representative trials** from the PPC array in monkey Mj, which followed a prior reward (PR, top) and a prior no-reward (PNR, bottom). The color maps show the phase gradient across the array for the 100ms prior to cue onset. The contour lines on the grid plot represent 8-Hz phase contours and the green arrows indicate the phase gradient vector at each location. The left polar plots illustrate the vector-sum of phase gradient vectors. **(B) Traveling waves on individual electrode arrays**. Across trial probability distribution of mean pre-cue wave strength values for -1.2s to 0s relative to cue onset, corresponding to dashed ROI in panel C, for PR and PNR trials. The horizontal bars represent mean and standard deviation. The mean pre-cue wave strength values of PR trials are significantly higher than PNR trials (rank-sum test PFC: Mj, p<0.05, Mc, p<0.01, PPC: Mj: p<0.01, Mc, p<0.01). **(C) Prior reward effects across time and frequency**. Average wave strength over PR (top) and PNR (bottom) trials calculated for 1.5Hz, 5ms windows of filtered LFP signals aligned to the reward cue onset. The dashed lines represent the cue onset (0s) and target onset (0.835 s); reward onset occurs 0.4 s after target acquisition. **(D) GLM coefficients for the prior trial effect**. Positive and negative coefficients respectively indicate enhancement and suppression of wave strength by a prior trial reward. The maps are thresholded to show only coefficients with significant p-values (p<0.05). Black contours show clusters of significant positive coefficients. A significant positive (red) region is present in the beta range, overlapping the pre-cue interval for all PFC and PPC arrays. Notably, there is no significant positive GLM coefficient around reward delivery (1.3s after cue onset), when licking is highest. **(E) Behavioral link with traveling wave strength is exclusive to reward history**. We examined the relation between traveling wave strength and expected reward, reward history, and reward outcome (see also Fig S4). Traces show the beta coefficients from the regression model indicating links between reward properties and wave strength at the ROI central frequency (PFC: Mc 22Hz, Mj 32Hz ; PPC: Mc 12Hz, Mj 10Hz). Shading represents the 95% confidence intervals of the coefficients. The reward history coefficients (red) are consistently non-zero for all PFC and PPC arrays in the pre-cue interval. **(F) Link between reward history and wave strength is not biased by oscillation power**. After statistically controlling for effects of LFP power, the residual wave strength was significantly higher for PR than PNR trials in both regions (all arrays p<0.01, ANOVA). Error bars represent SEM. **(G) Behavioral predictability of wave features**. Plot shows the coefficients from a multivariate GLM model that was trained to predict reward history based on three candidate features of traveling waves; wave strength, propagation direction (angular distance between wave direction and dominant propagation axis), and propagation speed. Traveling wave properties were measured for beta waves during the pre-cue interval. Black circles represent significant coefficients (p<0.05).

A similar pattern was present across the entire dataset, where TWs were reliably stronger on PR relative to PNR trials (Fig. 4B, top). The wave strength distribution was significantly greater on PR versus PNR trials in each array individually (non-parametric Wilcoxson rank-sum test dlPFC: Mj, p< 0.05, Mc, p<0.01, PPC: Mj: p<0.01, p<0.01). Although, as noted above, beta-band traveling waves were prominent throughout the pre-cue and post-cue (delay) epochs, the effect of the prior reward was limited to the pre-cue epoch, before the presentation of the cue signaling the current trial EV (Fig. 4C). This is seen by comparing the PR and PNR time-frequency maps (Fig. 4C) and was confirmed by a GLM analysis of the prior-trial effect (Fig. 4D).

The effect of the prior rewards was not explained by spurious factors related to licking, LFP power, or spatial sensitivity. The temporal dynamics of the prior reward modulations were clearly distinct from licking activity (which, unlike wave strength, peaked immediately before and after reward delivery) and there were no trial-by-trial correlations between licking and wave strength (Fig. S3). Second, the link between prior reward and traveling wave strength remained significant after accounting for LFP power (Fig. 4F; all arrays, p < 0.01). Finally, alpha-beta wave strength was not modulated by the locations of the cues or saccade targets, ruling out an explanation based on spatial factors (Fig. S4).

Importantly, additional analyses showed that the association we found was very specific between the prior reward (rather than other reward features) and the strength of the traveling waves (rather than other traveling wave features). Wave strength did not modulate immediately after reward delivery (Fig. S4), showing that the prior-reward modulations were not merely continuations of a response to the previous trial. Moreover, traveling waves did not modulate with the current trial’s EV (Fig. 4E, gray) or with the magnitude or probability predicted by the reward cue (Fig. S5). Finally, there was no consistent association between the prior reward and the speed and direction of the TW (Fig. 4G). These results reveal a very specific association between wave strength and prior rewards that does not extend to other behavioral measures or other aspects of the traveling waves.

### Traveling waves in the dlPFC encode the prior trial effect on the monkeys’ expectations

The analyses above show that the strength of the traveling waves at the start of a trial encodes the reward received on the previous trial. Can this pattern predict the animal’s memory for the prior reward? To examine this question, we used the monkeys’ licking response as a gauge of their sensitivity to the prior reward. Based on the distribution of licking rates on PR and PNR trials, we divided the trials into two groups, in which licking rates were consistent with the prior reward (high licking on PR trials and low licking on PNR trials) or inconsistent with that reward (low licking on PR trials and high licking on PNR trials; Fig. 5A, B). We reasoned that, if the effects of the prior reward on wave strength were independent of the monkeys’ memory for the previous reward, the effects should be equivalent on the two trial types. Conversely, if the wave strength effects were related to the monkeys’ memory for previous reward, they should be larger on trials when the monkeys retrieved the prior trial reward.

**Fig.5.**
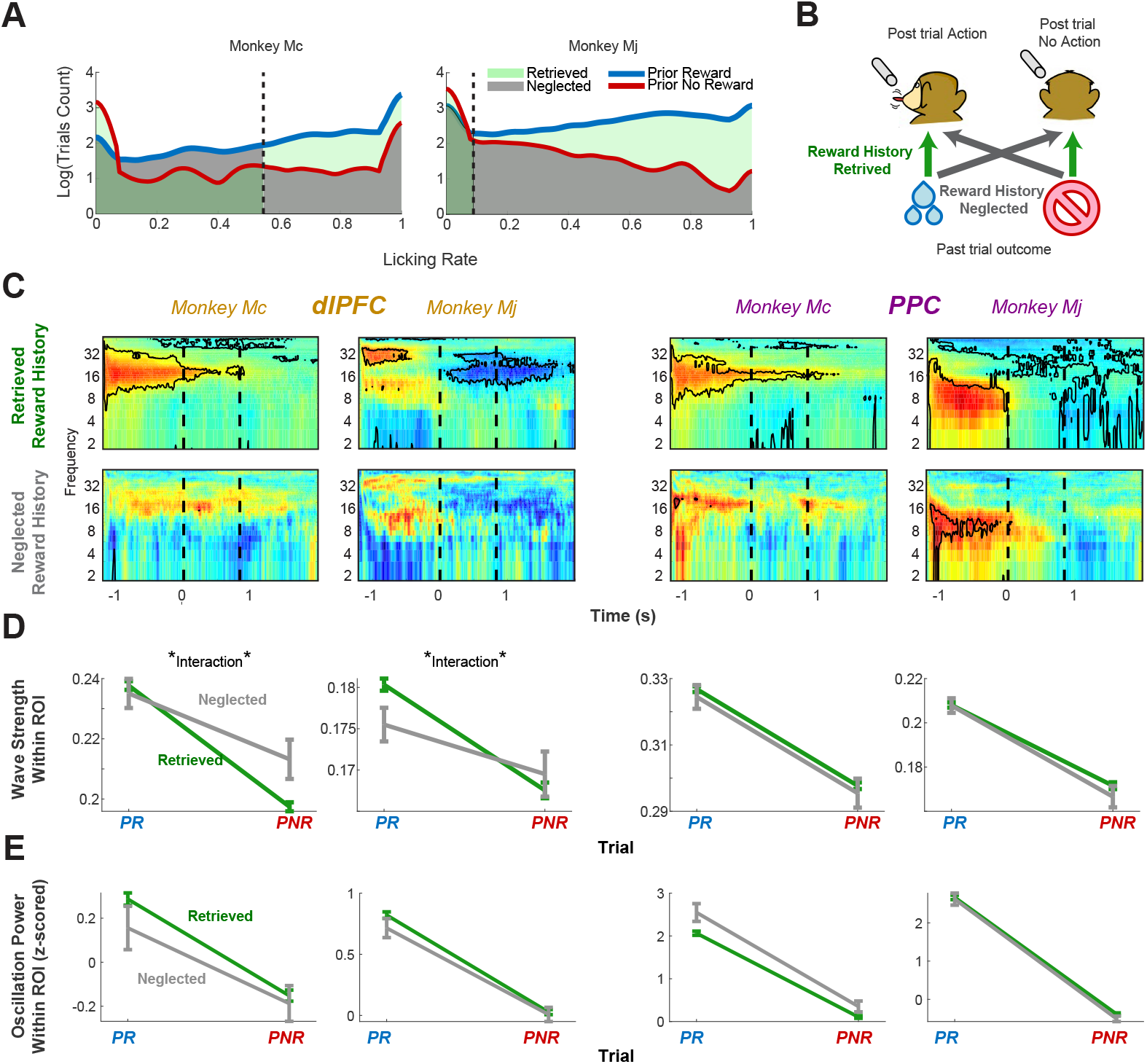
TWs in dlPFC are sensitive to the monkey’s memory of prior trial reward. **(A) Licking rates accurately categorize the prior trial reward**. The distributions of licking probability during the pre-cue epoch (500 ms to 0ms before cue onset), which differed between PR (blue) and PNR (red) trials in each monkey. We divided each distribution according to a decision criterion (vertical dashed line) that afforded maximal discrimination between PR and PNR trials (Methods). **(B) Illustration of the neglected and retrieved reward history trials**. The trials where the animal’s actions were consistent with the prior reward were considered as “retrieved reward history” (green) and, inversely, trails where the animal’s action was inconsistent with the prior reward were labeled “neglected reward history” (gray). **(C) Modulation of TW strength with prior reward is stronger when reward history is retrieved**. The top two panels represent the wave strength-PR regression coefficient map for each group of retrieved and neglected trials. The dashed lines represent cue (0s, left) and target (0.835s, right) onsets. **(D) dlPFC waves are more sensitive to monkey’s behavior**. Wave strength (y axis) is plotted as a function of prior reward (X-axis), and memory performance (Color). Data points represent the mean and SEM of averaged wave strength values within pre-cue positive ROIs. Significance determined from an analysis of variance. The interaction term for both frontal waves is significant (Mc p<0.03, Mj P<0.07) but not for the posterior waves (Mc p>0.9, Mj p>0.4). **(E) LFP power is not sensitive to memory performance**. Plot is analogous to panel D but for LFP power instead of TW strength. This result shows that LFP power does not change between neglected and retrieved power (p>0.4).

The effect of the prior rewards on traveling waves was sensitive to the monkeys’ behavior in the dlPFC but not in the PPC. In the dlPFC, the pre-cue GLM coefficients measuring the prior reward effect on the wave strength were significantly larger on consistent versus inconsistent trials (Fig. 5C; Mj and Mc: p<0.01, ttest), resulting in a significant interaction between trial type (PR, PNR) and behavior (consistent, inconsistent; ANOVA, interaction, Mj p < 0.07; Mc p < 0.03; Fig. 5D). In contrast, the PPC showed neither a significant difference between GLM coefficients (Fig. 5C) nor a significant interaction between trial type and behavior (Fig. 5D, right; Mj p >0.4, Mc p >0.9). A final ANOVA analysis with factors of area (dlPFC/PPC) and behavior (consistent, inconsistent) showed a significant area x behavior interaction (Mj < 0.05 Mc < 0.01). Interactions between prior reward and behavior were not found in the LFP power (Fig. 5E), ruling out that the effects on the wave strength were artifacts of the amplitude of LFP oscillations. Finally, the wave strength differences were replicated in an alternative analysis based on a median split of the correlation between licking and the prior reward (Fig. S5) ruling out that they were artefacts of a relatively small number of trials with inconsistent behavior. Thus, the strength of traveling waves in the frontal but not the parietal cortex increases when the animal has a strong memory of having recently received a reward.

## Discussion

We show that cortical TWs of beta-band oscillations are present in the monkey frontal and parietal cortex and convey information about recent rewards. In both areas, TW strength was enhanced by a recent reward and, in the dlPFC, this enhancement correlated with the effect of the reward on behavioral expectations. Thus, beyond their previously proposed roles in sensory and motor function, navigation, and memory (Davis et al., 2020; Lubenov and Siapas, 2009; Muller et al., 2018; Patel et al., 2012; Zhang et al., 2018), our results implicate TWs in reward processing in the fronto-parietal network.

Our finding that TWs were strongest in the beta frequency band (10–30 Hz) is consistent with a long line of work linking frontal beta-band oscillations with cognitive control and working memory (Babapoor-Farrokhran et al., 2017; Bhattacharya et al., 2022; Brincat and Miller, 2015; Miller et al., 2018; Womelsdorf, 2021). Studies have also suggested that beta-band oscillations behave as propagating TWs in sensory, motor and premotor structures (e.g., in humans ((Stolk et al., 2019; Takahashi et al., 2011), monkeys (Rubino et al., 2006; Takahashi et al., 2015), and cats (Roelfsema et al., 1997))). Our results extend the evidence for this claim to reward processing, and suggest that at least some of the previously reported effects of reward on the LFP oscillations in the frontal lobe (Taghizadeh et al., 2020) may in fact reflect signals that propagate in specific directions as planar traveling waves (Hamid et al., 2021).

Our finding that TWs are sensitive to rewards is consistent with the reward-selective spiking activity shown by many frontal and parietal cells (Foley et al., 2020; Kennerley et al., 2009; Taghizadeh et al., 2020; Wallis and Kennerley, 2010), and with the fact that these cells have sustained activity with relatively long time-constants and encode cross-trial memories, including about statistically irrelevant prior rewards (Abrahamyan et al., 2016; Akrami et al., 2018; Foley et al., 2020; Genovesio et al., 2014; Lee et al., 2012; Mansouri et al., 2006; Scott et al., 2017). However, our results suggest that TWs reflect unique aspects of reward computations that differ from those encoded in individual cells. First, we recently showed that, in the same behavioral task, individual dlPFC and 7a neurons responded much more to the delivery of reward relative to the prior trial reward (Foley et al., 2020; Kennerley et al., 2009; Wallis and Kennerley, 2010), a profile that dramatically differs from that of the TWs, which specifically encoded the prior reward but not the reward delivery or the expected reward. Second, our present result that both frontal and parietal TWs encoded the prior reward but only the frontal TWs encoded the effect of the reward on the monkeys’ anticipation, suggests a functional specialization that was not apparent in individual cells. Thus, while TW propagation across the entire fronto-parietal circuit may be important for maintaining reward memories, TWs specifically in the frontal lobe may gate the influence of these memories on immediate predictions and actions.

Our results also speak to the functional significance of the TW propagation direction. Because each cycle of a traveling wave is coupled to the activity of individual neurons (Bahramisharif et al., 2013; Buzsáki et al., 2012; Canolty et al., 2006; Katzner et al., 2009; Osipova et al., 2008) the presence of a spatially propagating TW suggests that a locus of neuronal spiking and/or subthreshold activity is moving across the cortex in a consistent direction (Swadlow and Alonso, 2009; Takahashi et al., 2015). However, the mechanisms and functional consequences of propagation direction are under debate. While some cognitive operations have been linked with distinctive propagation directions (Alamia and VanRullen, 2019; Friston, 2019; Pang et al., 2020), others seem to be related simply to the strength, but not the direction of TWs (Benucci et al., 2007; Davis et al., 2020; Nauhaus et al., 2009; Rubino et al., 2006; Swadlow and Alonso, 2009).

Our results combine elements of these views. On one hand, we find that prior rewards were associated with the strength but not specific direction of the TW. This result is consistent with the fact that reward properties are not known to be spatially mapped across the frontal or parietal cortex, and suggests that reward memory is associated with stronger but non-directional communication among reward-sensitive cells. On the other hand, rather than being uniformly distributed, TWs propagated primarily along two directions aligned with one spatial axis. This pattern may indicate preferential connectivity along anatomical axes orthogonal to a sulcus (Rubino et al., 2006) or, alternatively, may be due to the fact that our microarrays provided a narrow view, through a ∼10 mm^2^ aperture, into what may have been a more complex (e.g., spiral or circular) pattern over larger cortical areas (Bhattacharya et al., 2022; Ermentrout and Kleinfeld, 2001).

In sum, our findings show that TWs convey distinct aspects of reward computations relative to those conveyed by local LFP oscillations or individual cells, emphasizing the importance of understanding the spatial organization of LFP oscillations in multichannel recordings beyond measurements of individual sites (Muller et al., 2018).

## Methods

### Training and task

Two monkeys (Macaca mulatta), named Mc and Mj, performed a visually guided saccade task to obtain probabilistic rewards signaled by visual cues (Foley et al., 2020). Before neural recordings began, the monkeys were familiarized with 20 visual cues, signaling 7 levels of expected value (EV) through 10 distinct combinations of reward magnitude and probability. To perform the saccade task, the monkeys achieved central fixation, followed by the presentation of a randomly selected reward cue, a 600-ms memory period, and the target for the instructed saccade. After making the required saccade, the monkeys received one of two outcomes: reward omission or a reward with variable magnitude according to the contingencies signaled by the cue. A stereotyped tone marked the end of the post-saccadic hold period and the onset of the outcome epoch on reward and no-reward trials. The cue and target locations were independently randomized so that the cue was not predictive of the optimal action, and cues were equated for discriminability and counterbalanced across monkeys, with 2 cues assigned to each reward contingency. Together, these features of the task design allowed us to examine reward-related responses independently of visual confounds, learning or the planning of instrumental actions.

### Neural Recording

After completing behavioral training, each monkey was implanted with two 48-electrode Utah arrays (electrode length 1.5 mm) arranged in rectangular grids (monkey Mc, 7 × 7 grid; monkey Mj, 5 × 10 grid) and positioned in the pre-arcuate portion of the dlPFC and the posterior portion of area 7A. Data were recorded using the Cereplex System (Blackrock, Salt Lake City, Utah) over 18 sessions spanning 4 months after array implantation in monkey Mj, and 12 sessions spanning 2 months after implantation in monkey Mc.

### Spectral analysis of the LFP

The raw LFP from each electrode and trial were low pass filtered at 200 Hz, and notch filtered at 60 Hz to remove line noise. We removed trials with artifacts, such as outlier step-like artifacts in LFP traces, by identifying the peak-to-peak amplitude of the broadband LFP trace in each trial. We removed any trials for which this z transformed measure was more than half a standard deviation away from the mean across all trials.The dominant oscillation frequency is calculated based on the multitaper power spectral density estimation (Babadi and Brown, 2014) implemented with Chronux (Bokil et al., 2010). Power spectra were computed separately for each channel and trial and the background power spectrum was removed through 1/f slope estimation (Kasdin, 1995).

To identify the power of oscillations at different frequencies throughout the task, the spectrograms of were computed using the continuous wavelet transform (CWT) with analytic Morse wavelet family scales (Muthukumaraswamy and Johnson, 2004) corresponding to wavelet center frequencies of 1 to 60 Hz. The hyperparameters of the wavelet transform were optimized automatically based on wavelet’s energy spread [gamma = 3, voice per octave = 10, time-bandwidth = 60, and signal sampling rate = 200 Hz]. For follow-up analyses at specific frequencies, we applied a zero-phase lag Butterworth second-order filter with a bandwidth of 1.5 Hz, which prevents relative phase deformation.

### Characterizing traveling waves

We implemented a version of the traveling-wave characterization method from Zhang et al. (Zhang et al., 2018). As is shown in Fig. 3A, the spectrum of recorded LFPs often contains multimodal peaks in the frequency domain, so we systematically applied our analyses at a broad range of frequencies. As is illustrated in Fig. S1, at frequencies ranging from 2 to 50 Hz, we first applied a zero-phase band-pass filter with a fixed width of ±1.5Hz. We used this approach to extract the signals at each electrode, and then applied the Hilbert transform to the filtered voltage signal, V(x,y,t), recorded by an electrode located at position x,y on the array.

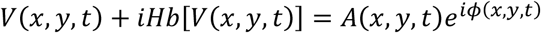

where *Hb* is the Hilbert transform operator and A(x,y,t) is the instantaneous recorded voltage, and *ϕ*(*x,y,t*)is the instantaneous phase. This procedure extracts a separate phase for each electrode at each time point, which we then used to calculate spatial phase gradients throughout the task for each recording site.

We measured the phase gradient at each electrode and timepoint by fitting a plane to the measured phase values across the grid. This regression fit a plane that predicted the phase at each electrode, with the non-zero slopes *k*_*x*_ and *k*_*y*_ indicating that there is a systematic phase gradient across the grid.

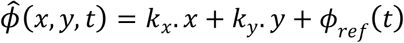

This model thus fits the relation between the measured phase gradient and the electrode position (Fig. S1). We measured this goodness of fit quantitatively by measuring the Phase Gradient Directionality (PGD) (Rubino et al., 2006), which is the Pearson correlation between the actual phase of traveling wave and the predicted phase from the best-fitting planar wave model at each time point (*t*):

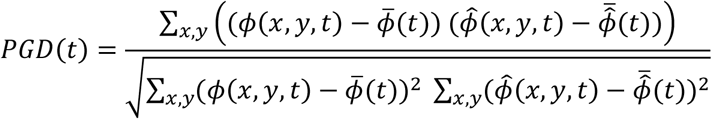

Here,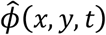 is the predicted phase of the signal at each location and timepoint, and *ϕ*_*ref*_(*t*)is the phase on the reference electrode. Then, using the slope of the fitted plane wave, (*k*_*x*_ and *k*_*y*_)we identified propagation direction as below:

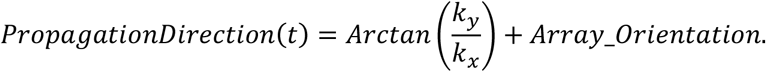

Finally, we measured the spatial propagation speed, *ν*, following the equation for measuring waves speed in physical systems (Elmore et al., 1985).

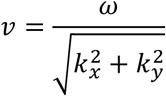

### GLM model

To measure the effect of task parameters on wave strength (PGD) and oscillation power, we implemented GLM models, which produced a time-frequency map of coefficients showing the relations between task parameters, PGD, and LFP Power for each electrode (Fig. 4 & Fig S.4). We then identified significant regions of Interest (ROI) within the GLM coefficient maps, which were specific time–frequency windows for which the beta coefficients were significantly non-zero (F-test with p<0.05).

### Residual Wave Strength

Our early observations indicate that TW strength was enhanced in the frequency bands that showed peaks in the LFP power spectrum. Given that these frequencies show both TWs and large-amplitude oscillations, we used a statistical approach to test which was more closely related to the behavior, the TW specifically or, more generally, the power of the underlying oscillation. First, for each trial, we computed the average oscillation power and TW strength (Fig 4D). We then verified that our behavioral links with TWs were not driven by effects of power by using a linear regression to model the relation between TWs strength and power. By creating a trial-level prediction of TW strength from power, the residual from this regression provided a measure of TW strength that was not predicted by power, independently for each array. We then reran our initial analyses instead with this power-independent TWs strength and found that it was significantly higher for PR trials compared to PNR trials in each recording grid (all arrays P<0.01, ANOVA). This approach thus confirmed that our main results reflected TW strength specifically, rather than power.

### ROC analysis

We used a Receiver Operating Characteristic (ROC) curve to quantify the distinguishability of a prior trial’s outcome given the monkeys’ behavior in the following trial. We took the licking rate from [0 to - 500ms] before cue onset for each trial and generated a ROC curve to predict whether a reward had been previously released, as a function of different decision criteria (i.e., licking rates as threshold). The Area Under the Curve (AUC) was high for both monkeys [Mc AUC = 0.85; Mj AUC = 0.89], indicating that licking behavior can reliably predict the precedent trial’s reward outcome. For each monkey, the optimal decision criteria were defined based on the maximum distance to the diagonal line in the ROC curve. Our labels of neglected and retrieved trials were selected based on this optimal decision criterion.

## Acknowledgments

We acknowledge support from Columbia University’s Research Initiatives in Science & Engineering program.

## Supplemental Materials

**Video S1:**
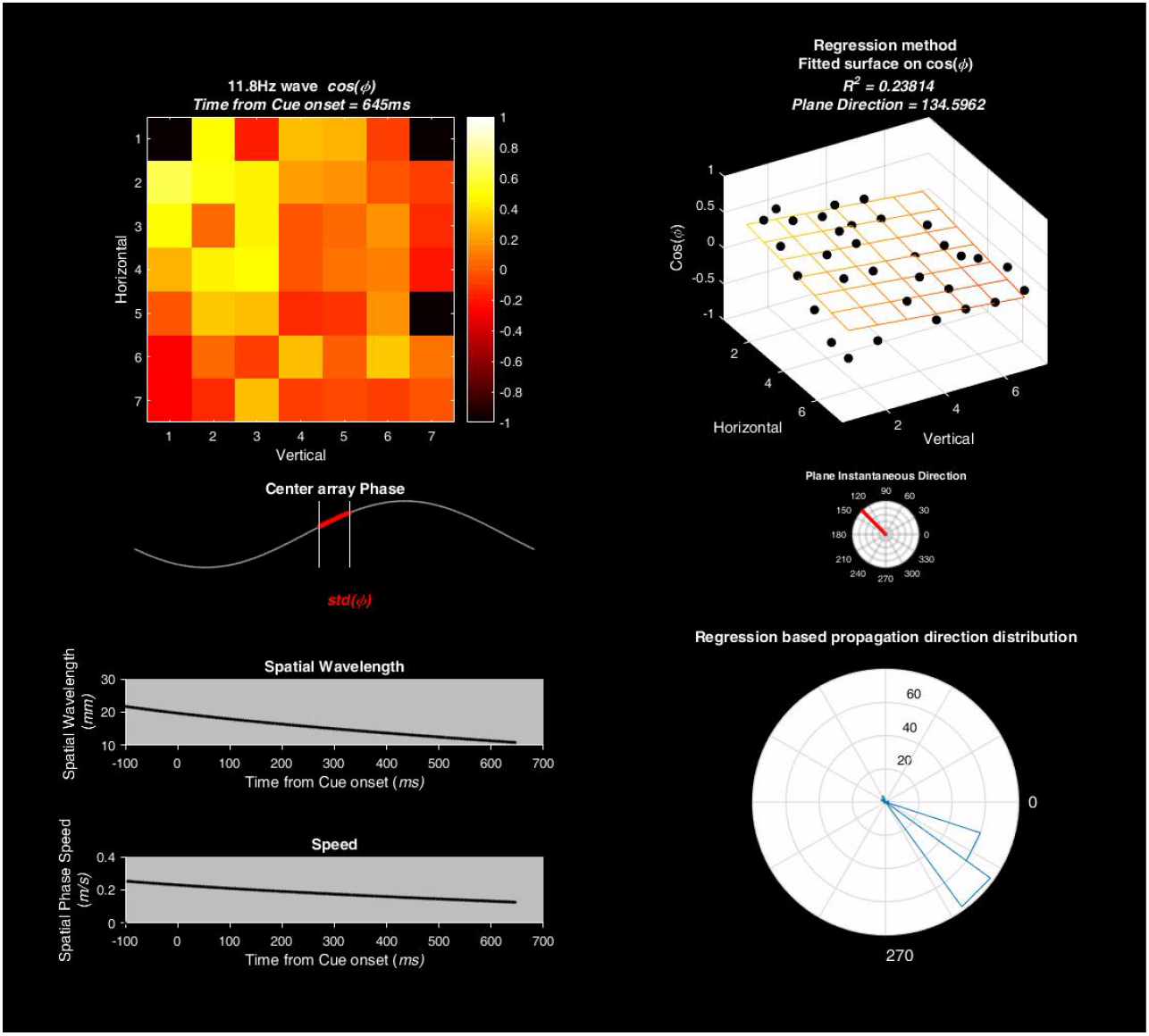
Example video showing one representative TW: [One frame attached below] Data show an example trial (recording session 3, trial 175) from monkey Mj’s array PPC. Animation includes the phase values of an 11.8-Hz oscillation and the illustration of fitting a traveling wave on the spatial phase distribution. The instantaneous properties of the wave, including speed and direction, are also illustrated. The single red arrow represents the mean direction across electrodes.

**Fig. S1.**
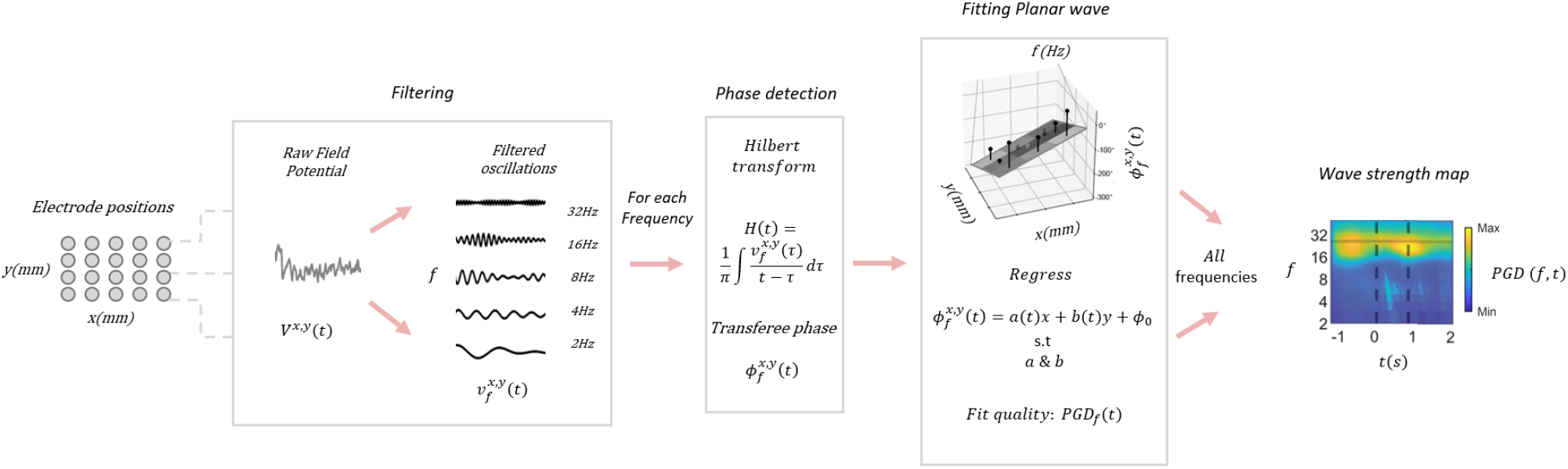
Wave strength map calculation. Flowchart illustrating the analysis paradigm for obtaining PGD maps from the raw data. For each recording electrode in position x and y, the raw LFP signal V^x,y^(t) decomposed to oscillatory components v^x,y^_f_ (t) with a frequency range f from 2 to 50 Hz. For each decomposed oscillation the phase of oscillation is extracted using the Hilbert transform and then all the electrode phases are used to measure the relation between phase and location. The traveling wave properties for each frequency were calculated independently and then merged for display in the wave strength map.

**Fig. S2.**
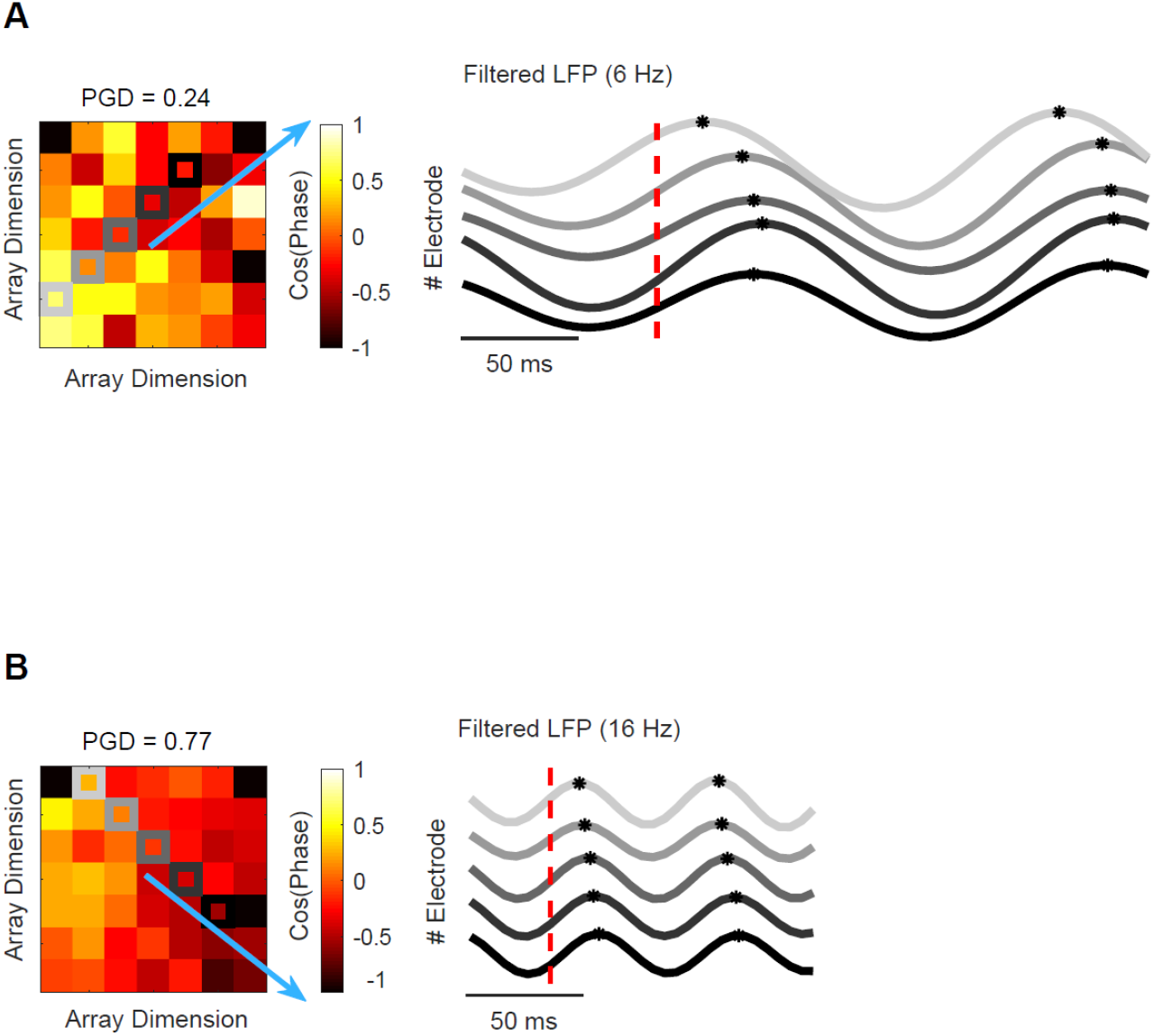
Example of high and low frequency traveling waves. **(A) Slow oscillations as Traveling Waves**. Representative trial from monkey Mj’s posterior array showing low-frequency oscillation (filtered at 6Hz) across the highlighted adjacent electrodes (light to dark gray) within the pre-cue interval. Black stars indicate oscillation peaks in each electrode, showing a systematic phase shift across adjacent electrodes. The left phase map illustrates the spatial distribution of phases across the recording array at a single time snapshot, indicated with the dashed red line in the right panel. The blue arrow represents the overall direction of phase propagation. **(B) Fast oscillation as Traveling Waves**. Illustration of a representative traveling wave with a high frequency (filtered at 16 Hz) from the same array.

**Fig. S3.**
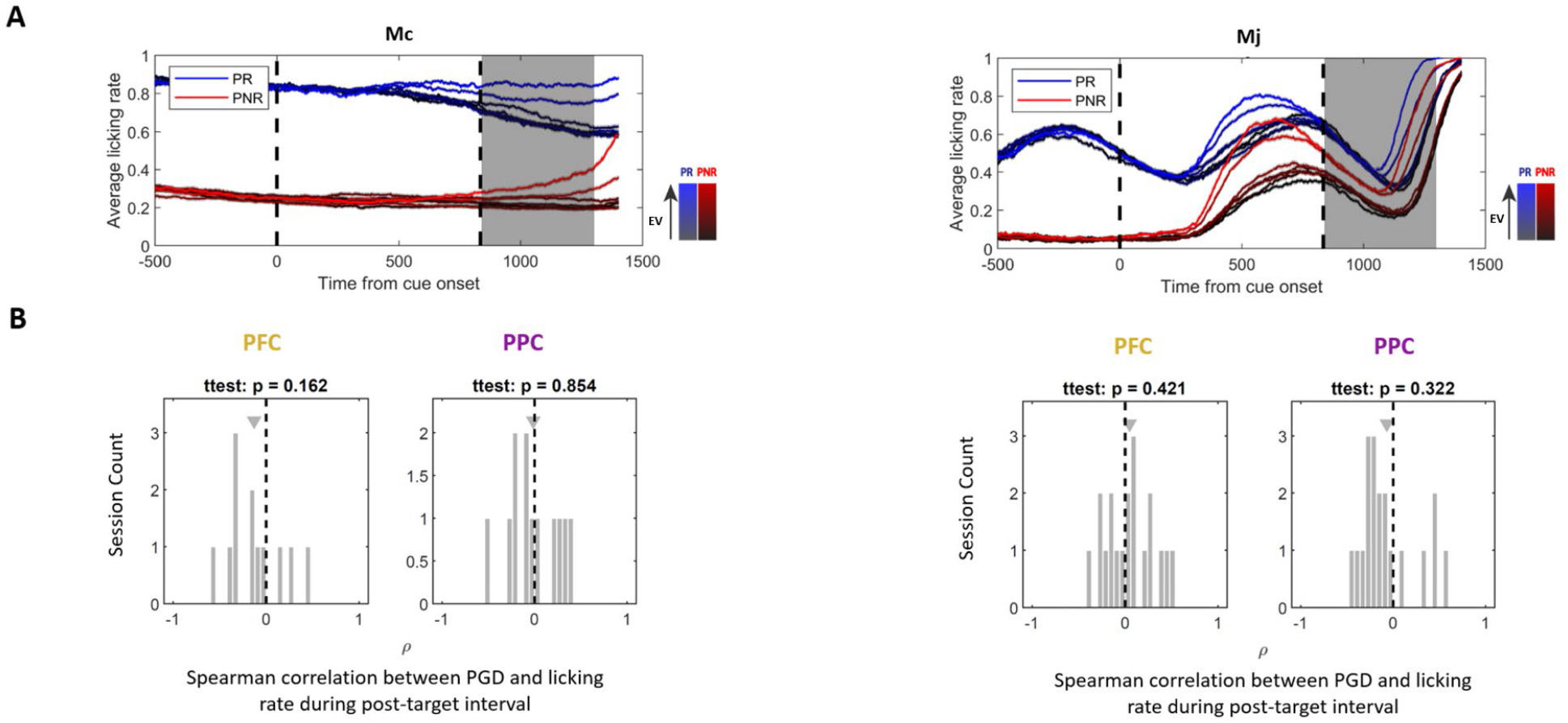
PGD is not an artifact of licking activity. **(A) Licking rate dynamics**. Top panels represent the temporal dynamics of licking rate, separately computed for PR and PNR trials with different expected values (stronger colors indicate lower EV). **(B) Analysis indicating that there is no correlation between licking and wave strength around reward delivery**. The average licking rate within the shaded interval (500ms post-target interval) was z-scored based on PR condition and then binned by the expected value on each trial. Horizontal axis indicates the spearman correlation between wave strength (measured by PGD) and the z-scored licking rate in the same interval, for each session. Plot indicates a histogram of the spearman correlation coefficient. None of the distributions differ from zero (p’s>0.1), indicating that wave strength fluctuations are independent of licking rate in this time interval.

**Fig. S4.**
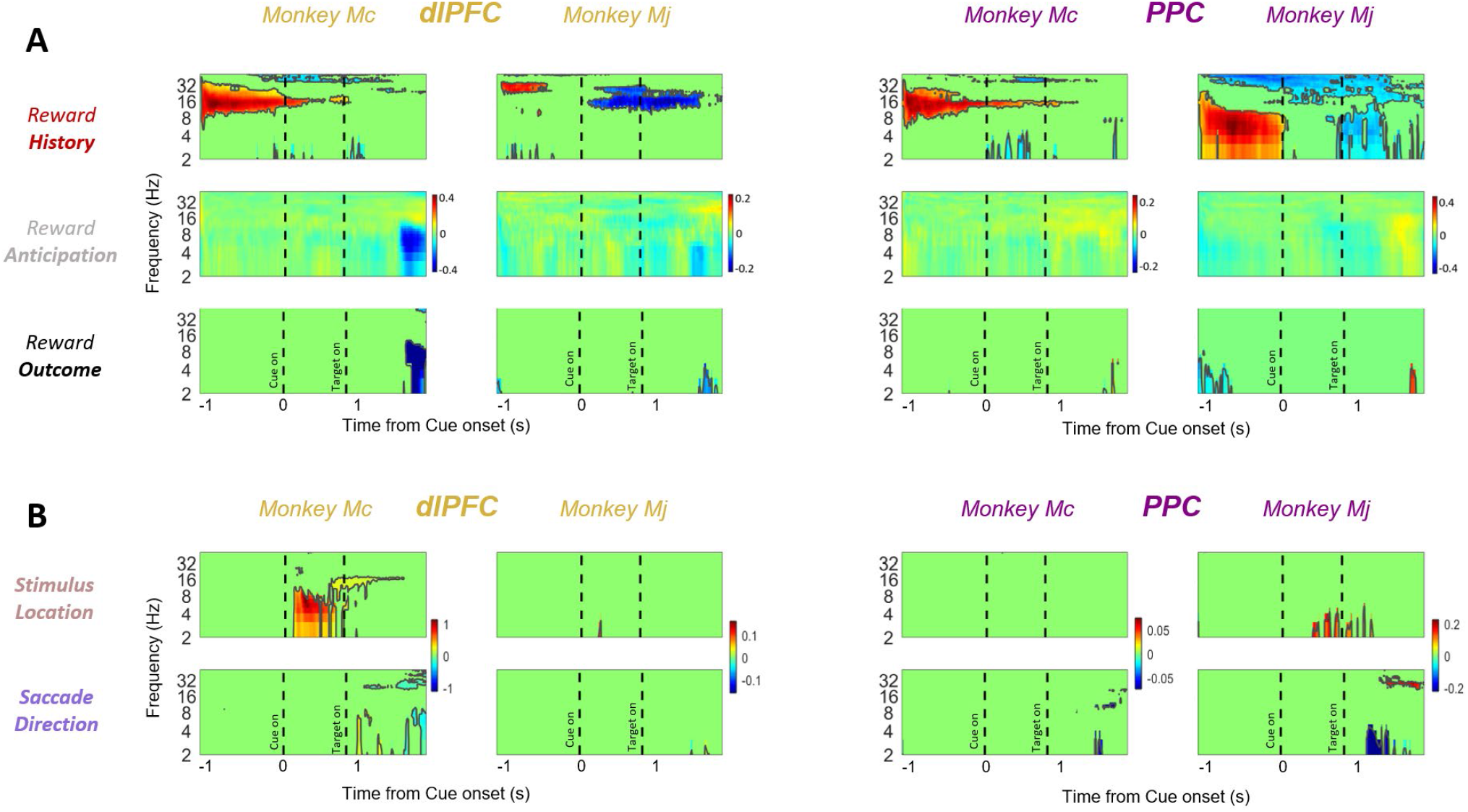
Behavioral relevance of traveling wave strength. **(A) Model coefficients showing link between TW strength and reward parameters**. Each panel shows the result of a linear model relating traveling wave strength with different reward parameters, across time and frequency map. Top: Prior Reward (Reward History), Middle: Expected Value of Reward (Reward Anticipation), bottom: Current Reward (Reward Outcome). The plotted values are normalized for each array based on the maximum across all parameters for each array. Black line indicates clusters of significance at p<0.05. The dashed lines represent the times of cue onset (0s) and target onset (0.835s). **(B) Model coefficients relating TW strength to visuomotor behaviors**. Overall, although some visually evoked patterns are low frequencies are present, they are only a weak trend and occur at different timepoints and frequencies compared to the reward effects thus indicating that they are a distinct phenomenon.

**Fig. S5.**
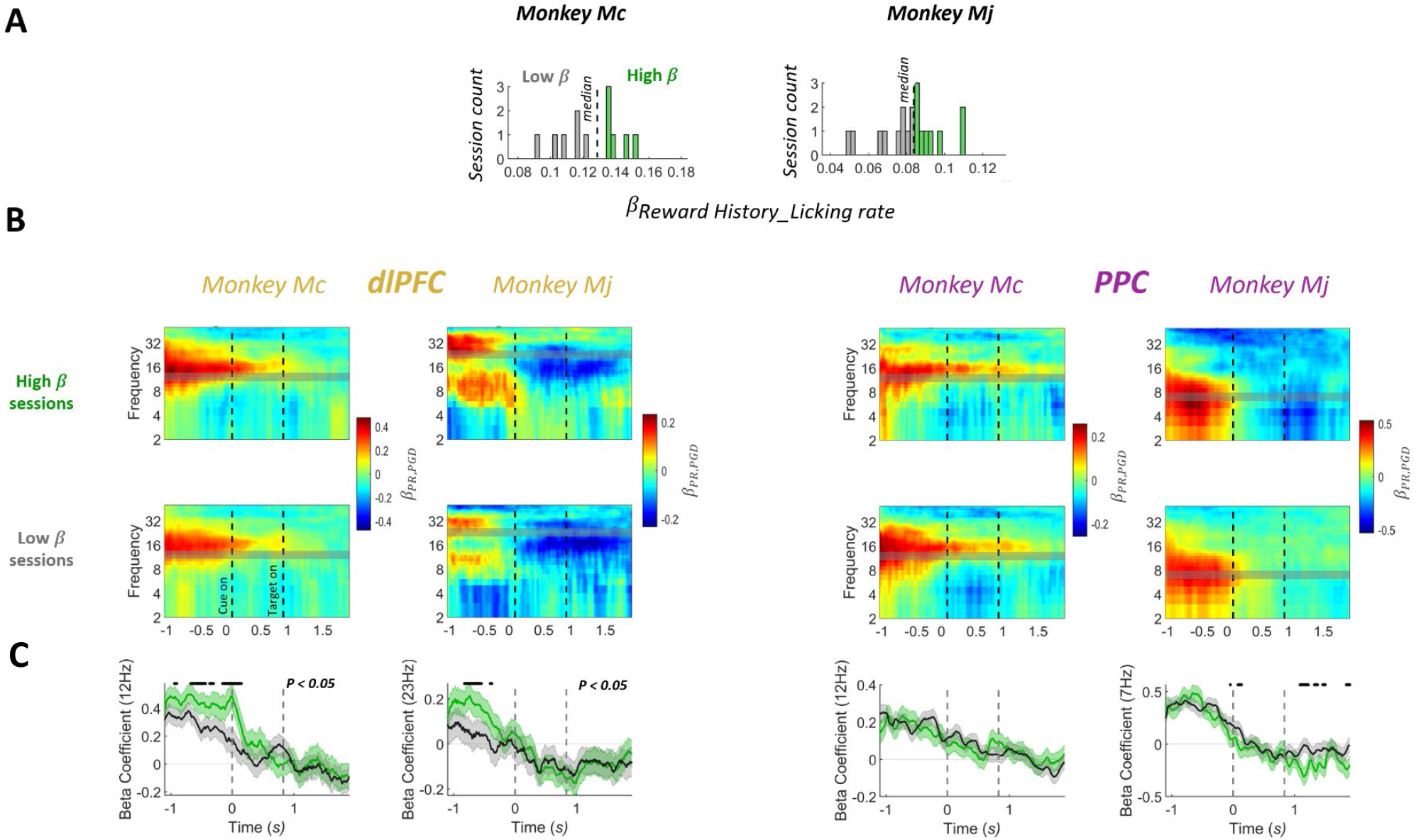
Replication of figure 5, with data split between sessions that show high and low relation between licking and past reward. **(A) Methodology for categorize sessions based on the animal’s behavioral sensitivity to prior reward**. The high (green) and low (gray) behavior-dependent sessions have been labeled based on a median split of their coefficient. **(B) Beta coefficient maps showing the link between wave strength and PR separately for each of high and low behaviorally sensitive sessions. (C) Plots indicating detail on the effects at each animal’s peak frequency**. Shaded error bars represent SEM. Here, the green trace is higher than the gray trace (p <0.05, t-test) during the pre-cue interval for dlPFC arrays, which is consistent with our interpretation that the strength of TWs in the dlPFC are consistently related to the monkey’s memory for the prior trial’s reward.

## References

1. Abrahamyan, A., Silva, L.L., Dakin, S.C., Carandini, M., and Gardner, J.L. (2016). Adaptable history biases in human perceptual decisions. Proceedings of the National Academy of numSciences 113, E3548–E3557.

2. Akrami, A., Kopec, C.D., Diamond, M.E., and Brody, C.D. (2018). Posterior parietal cortex represents sensory history and mediates its effects on behaviour. Nature 554, 368–372.

3. Alamia, A., and VanRullen, R. (2019). Alpha oscillations and traveling waves: Signatures of predictive coding? PLoS Biol. 17, e3000487.

4. Alamia, A., Timmermann, C., Nutt, D.J., VanRullen, R., and Carhart-Harris, R.L. (2020). Correction: DMT alters cortical travelling waves. Elife 9.

5. Babadi, B., and Brown, E.N. (2014). A review of multitaper spectral analysis. IEEE Transactions on Biomedical Engineering 61, 1555–1564.

6. Babapoor-Farrokhran, S., Vinck, M., Womelsdorf, T., and Everling, S. (2017). Theta and beta synchrony coordinate frontal eye fields and anterior cingulate cortex during sensorimotor mapping. Nat. Commun. 8, 13967.

7. Bahramisharif, A., van Gerven, M.A.J., Aarnoutse, E.J., Mercier, M.R., Schwartz, T.H., Foxe, J.J., Ramsey, N.F., and Jensen, O. (2013). Propagating neocortical gamma bursts are coordinated by traveling alpha waves. J. Neurosci. 33, 18849–18854.

8. Bakker, R., Tiesinga, P., and Kötter, R. (2015). The scalable brain atlas: instant web-based access to public brain atlases and related content. Neuroinformatics 13, 353–366.

9. Balasubramanian, K., Papadourakis, V., Liang, W., Takahashi, K., Best, M.D., Suminski, A.J., and Hatsopoulos, N.G. (2020). Propagating motor cortical dynamics facilitate movement initiation. Neuron 106, 526–536.

10. Benucci, A., Frazor, R.A., and Carandini, M. (2007). Standing waves and traveling waves distinguish two circuits in visual cortex. Neuron 55, 103–117.

11. Bhattacharya, S., Brincat, S.L., Lundqvist, M., and Miller, E.K. (2022). Traveling waves in the prefrontal cortex during working memory. PLoS Comput. Biol. 18, e1009827.

12. Bokil, H., Andrews, P., Kulkarni, J.E., Mehta, S., and Mitra, P.P. (2010). Chronux: a platform for analyzing neural signals. J. Neurosci. Methods 192, 146–151.

13. Brincat, S.L., and Miller, E.K. (2015). Frequency-specific hippocampal-prefrontal interactions during associative learning. Nat. Neurosci. 18, 576–581.

14. Buzsáki, G., Anastassiou, C.A., and Koch, C. (2012). The origin of extracellular fields and currents—EEG, ECoG, LFP and spikes. Nat. Rev. Neurosci. 13, 407–420.

15. Calabrese, E., Badea, A., Coe, C.L., Lubach, G.R., Shi, Y., Styner, M.A., and Johnson, G.A. (2015). A diffusion tensor MRI atlas of the postmortem rhesus macaque brain. Neuroimage 117, 408–416.

16. Canolty, R.T., Edwards, E., Dalal, S.S., Soltani, M., Nagarajan, S.S., Kirsch, H.E., Berger, M.S., Barbaro, N.M., and Knight, R.T. (2006). High Gamma Power Is Phase-Locked to Theta Oscillations in Human Neocortex. Science.

17. Davis, Z.W., Muller, L., Martinez-Trujillo, J., Sejnowski, T., and Reynolds, J.H. (2020). Spontaneous travelling cortical waves gate perception in behaving primates. Nature 587, 432–436.

18. Davis, Z.W., Benigno, G.B., Fletterman, C., Desbordes, T., Steward, C., Sejnowski, T.J., H Reynolds, J., and Muller, L. (2021). Spontaneous traveling waves naturally emerge from horizontal fiber time delays and travel through locally asynchronous-irregular states. Nat. Commun. 12, 6057.

19. Dickey, C.W., Sargsyan, A., Madsen, J.R., Eskandar, E.N., Cash, S.S., and Halgren, E. (2021). Travelling spindles create necessary conditions for spike-timing-dependent plasticity in humans. Nat. Commun. 12, 1027.

20. Elmore, W.C., Elmore, W.C., and Heald, M.A. (1985). Physics of waves (Courier Corporation).

21. Ermentrout, G.B., and Kleinfeld, D. (2001). Traveling electrical waves in cortex: insights from phase dynamics and speculation on a computational role. Neuron 29, 33–44.

22. Fell, J., and Axmacher, N. (2011). The role of phase synchronization in memory processes. Nat. Rev. Neurosci. 12, 105–118.

23. Foley, N.C., Cohanpour, M., Semework, M., Sheth, S.A., and Gottlieb, J. (2020). Population coding of reward prediction errors through opponent organization in the fronto parietal network.

24. Fries, P. (2005). A mechanism for cognitive dynamics: neuronal communication through neuronal coherence. Trends Cogn. Sci. 9, 474–480.

25. Friston, K.J. (2019). Waves of prediction. PLoS Biol. 17, e3000426.

26. Genovesio, A., Tsujimoto, S., Navarra, G., Falcone, R., and Wise, S.P. (2014). Autonomous encoding of irrelevant goals and outcomes by prefrontal cortex neurons. J. Neurosci. 34, 1970–1978.

27. Girard, P., Hupé, J.M., and Bullier, J. (2001). Feedforward and feedback connections between areas V1 and V2 of the monkey have similar rapid conduction velocities. J. Neurophysiol. 85, 1328–1331.

28. González-Burgos, G., Barrionuevo, G., and Lewis, D.A. (2000). Horizontal synaptic connections in monkey prefrontal cortex: an in vitro electrophysiological study. Cereb. Cortex 10, 82–92.

29. Halgren, M., Ulbert, I., Bastuji, H., Fabó, D., Erőss, L., Rey, M., Devinsky, O., Doyle, W.K., Mak-McCully, R., Halgren, E., et al. (2019). The generation and propagation of the human alpha rhythm. Proceedings of the National Academy of Sciences 116, 23772–23782.

30. Hamid, A.A., Frank, M.J., and Moore, C.I. (2021). Wave-like dopamine dynamics as a mechanism for spatiotemporal credit assignment. Cell 184, 2733–2749.e16.

31. Kasdin, N.J. (1995). Discrete simulation of colored noise and stochastic processes and 1/f/sup/spl alpha//power law noise generation. Proc. IEEE 83, 802–827.

32. Katzner, S., Nauhaus, I., Benucci, A., Bonin, V., Ringach, D.L., and Carandini, M. (2009). Local origin of field potentials in visual cortex. Neuron 61, 35–41.

33. Kennerley, S.W., Dahmubed, A.F., Lara, A.H., and Wallis, J.D. (2009). Neurons in the frontal lobe encode the value of multiple decision variables. J. Cogn. Neurosci. 21, 1162–1178.

34. Kleen, J.K., Chung, J.E., Sellers, K.K., Zhou, J., Triplett, M., Lee, K., Tooker, A., Haque, R., and Chang, E.F. (2021). Bidirectional propagation of low frequency oscillations over the human hippocampal surface. Nat. Commun. 12, 1–10.

35. Lak, A., Hueske, E., Hirokawa, J., Masset, P., Ott, T., Urai, A.E., Donner, T.H., Carandini, M., Tonegawa, S., Uchida, N., et al. (2020). Reinforcement biases subsequent perceptual decisions when confidence is low, a widespread behavioral phenomenon. Elife 9, e49834.

36. Lee, D., Seo, H., and Jung, M.W. (2012). Neural basis of reinforcement learning and decision making. Annu. Rev. Neurosci. 35, 287–308.

37. Lubenov, E.V., and Siapas, A.G. (2009). Hippocampal theta oscillations are travelling waves. Nature 459, 534–539.

38. Mansouri, F.A., Matsumoto, K., and Tanaka, K. (2006). Prefrontal cell activities related to monkeys’ success and failure in adapting to rule changes in a Wisconsin Card Sorting Test analog. J. Neurosci. 26, 2745–2756.

39. Massimini, M., Huber, R., Ferrarelli, F., Hill, S., and Tononi, G. (2004). The sleep slow oscillation as a traveling wave. Journal of Neuroscience 24, 6862–6870.

40. Miller, E.K., Lundqvist, M., and Bastos, A.M. (2018). Working Memory 2.0. Neuron 100, 463–475.

41. Muller, L., Piantoni, G., Koller, D., Cash, S.S., Halgren, E., and Sejnowski, T.J. (2016). Rotating waves during human sleep spindles organize global patterns of activity that repeat precisely through the night. Elife 5, e17267.

42. Muller, L., Chavane, F., Reynolds, J., and Sejnowski, T.J. (2018). Cortical travelling waves: mechanisms and computational principles. Nat. Rev. Neurosci. 19, 255.

43. Muthukumaraswamy, S.D., and Johnson, B.W. (2004). Primary motor cortex activation during action observation revealed by wavelet analysis of the EEG. Clin. Neurophysiol. 115, 1760–1766.

44. Nauhaus, I., Busse, L., Carandini, M., and Ringach, D.L. (2009). Stimulus contrast modulates functional connectivity in visual cortex. Nat. Neurosci. 12, 70.

45. Osipova, D., Hermes, D., and Jensen, O. (2008). Gamma power is phase-locked to posterior alpha activity. PLoS One 3, e3990.

46. Pang, Z., Alamia, A., and VanRullen, R. (2020). Turning the Stimulus On and Off Changes the Direction of α Traveling Waves. eNeuro 7.

47. Patel, J., Fujisawa, S., Berényi, A., Royer, S., and Buzsáki, G. (2012). Traveling theta waves along the entire septotemporal axis of the hippocampus. Neuron 75, 410–417.

48. Roelfsema, P.R., Engel, A.K., König, P., and Singer, W. (1997). Visuomotor integration is associated with zero time-lag synchronization among cortical areas. Nature 385, 157–161.

49. Rubino, D., Robbins, K.A., and Hatsopoulos, N.G. (2006). Propagating waves mediate information transfer in the motor cortex. Nat. Neurosci. 9, 1549.

50. Sato, T.K., Nauhaus, I., and Carandini, M. (2012). Traveling waves in visual cortex. Neuron 75, 218–229.

51. Scott, B.B., Constantinople, C.M., Akrami, A., Hanks, T.D., Brody, C.D., and Tank, D.W. (2017). Fronto-parietal cortical circuits encode accumulated evidence with a diversity of timescales. Neuron 95, 385–398.

52. Stolk, A., Brinkman, L., Vansteensel, M.J., Aarnoutse, E., Leijten, F.S.S., Dijkerman, C.H., Knight, R.T., de Lange, F.P., and Toni, I. (2019). Electrocorticographic dissociation of alpha and beta rhythmic activity in the human sensorimotor system. Elife 8, e48065.

53. Swadlow, H.A., and Alonso, J.-M. (2009). Spikes are making waves in the visual cortex. Nat. Neurosci. 12, 10–11.

54. Taghizadeh, B., Foley, N.C., Karimimehr, S., Cohanpour, M., Semework, M., Sheth, S.A., Lashgari, R., and Gottlieb, J. (2020). Reward uncertainty asymmetrically affects information transmission within the monkey fronto-parietal network. Communications Biology 3, 1–11.

55. Takahashi, K., Saleh, M., Penn, R.D., and Hatsopoulos, N.G. (2011). Propagating waves in human motor cortex. Front. Hum. Neurosci. 5, 40.

56. Takahashi, K., Kim, S., Coleman, T.P., Brown, K.A., Suminski, A.J., Best, M.D., and Hatsopoulos, N.G. (2015). Large-scale spatiotemporal spike patterning consistent with wave propagation in motor cortex. Nat. Commun. 6, 1–11.

57. Wallis, J.D., and Kennerley, S.W. (2010). Heterogeneous reward signals in prefrontal cortex. Curr. Opin. Neurobiol. 20, 191–198.

58. Womelsdorf, T. (2021). Translating Expectation into Visual Selection through a Beta-Synchronous Fronto-Parietal Neural Subnetwork. Neuron 109, 8–10.

59. Zanos, T.P., Mineault, P.J., Nasiotis, K.T., Guitton, D., and Pack, C.C. (2015). A sensorimotor role for traveling waves in primate visual cortex. Neuron 85, 615–627.

60. Zhang, H., and Jacobs, J. (2015). Traveling theta waves in the human hippocampus. Journal of Neuroscience 35, 12477–12487.

61. Zhang, H., Watrous, A.J., Patel, A., and Jacobs, J. (2018). Theta and alpha oscillations are traveling waves in the human neocortex. Neuron 98, 1269–1281.

